# Ultra-sparse connectivity within the lateral hypothalamus

**DOI:** 10.1101/2020.04.25.061564

**Authors:** Denis Burdakov, Mahesh M. Karnani

## Abstract

The lateral hypothalamus (LH) contains neuronal populations which generate fundamental behavioural actions such as feeding, sleep, movement, attack and evasion. Their activity is also correlated with various appetitive and consummatory behaviours as well as reward seeking. It is unknown how neural activity within and among these populations is coordinated. One hypothesis postulates that they communicate using inhibitory and excitatory synapses, forming local microcircuits. We inspected this hypothesis using quadruple whole cell recordings and optogenetics to screen thousands of potential connections in brain slices. In contrast to the neocortex, we found near zero connectivity within the LH. In line with its ultra-sparse intrinsic connectivity, we found that the LH does not generate local beta and gamma oscillations. This suggests that LH neurons integrate incoming input within individual neurons rather than through local network interactions, and that input from other brain structures is decisive for selecting active populations in LH.

## Introduction

The lateral hypothalamic area (LH) is a vital controller of arousal, feeding and metabolism (Bernardis and Bellinger, 1996; Bonnavion et al., 2016). It integrates external and internal sensory information and outputs to most of the brain as well as the autonomic nervous system. Synaptic efferents from many sources including the striatum and midbrain impinge on LH neurons through the medial forebrain bundle (Iyer et al., 2018), but it is unknown how these inputs are processed by LH synaptic networks. While the sensory and whole-body output properties of LH cell populations have received much interest, their intrinsic synaptic organization has remained largely unstudied.

It would make sense for LH neurons to be synaptically connected to each other, forming inhibitory and excitatory connections, as these microcircuits could help integrate and filter sensory information and activate populations as a unit (Karnani et al., 2020). Furthermore, they could allow coordinating activity between LH cell types, some of which have mutually exclusive behavioural effects, such as LH glutamatergic vglut2 and GABAergic VGAT neurons (Jennings et al., 2013, 2015; Li et al., 2018; Stamatakis et al., 2016), and orexin (orx) and melanin concentrating hormone (MCH) neurons (Adamantidis et al., 2007; Jego et al., 2013; Konadhode et al., 2013). Mutual inhibitory coordination of such behavioural control nodes has been a long-standing hypothesis (Tinbergen, 1951). However, classical Golgi staining studies did not find interneurons with locally ramifying axons in the LH (Millhouse, 1979, 1969) and recent studies have demonstrated a lack of local synaptic connectivity in neighboring subthalamic and thalamic areas (Hou et al., 2016; Steiner et al., 2019).

Studies with optogenetic circuit mapping within the LH have demonstrated only a minority of connections when a large pool of presynaptic neurons were activated (Apergis-Schoute et al., 2015; Ferrari et al., 2018; Kosse and Burdakov, 2019; Kosse et al., 2017). However, multiple patch clamp has not been used to study LH connectivity, aside from a limited dataset of MCH neurons where no connections were discovered (Apergis-Schoute et al., 2015). Therefore, we used quadruple whole cell recordings in acute brain slices to screen connectivity within the LH with standard methodology we previously used in neocortex (Jackson et al., 2018; Karnani et al., 2016a, 2016b). Finding a lack of local connectivity, we used optogenetic circuit mapping to study the strength of LH optogenetic responses and network oscillations, which we found to be consistent with ultra-sparse intrinsic connectivity within the LH.

## Results

### Ultra-sparse connectivity with multiple patch clamp recordings

To begin looking for synaptic connectivity within LH, we used standard methodology to cut brain slices and obtain multiple simultaneous whole cell patch clamp recordings in wild type animals (Figure 1A,B). We sequentially imposed a 50 Hz train of 5 action potentials on each recorded neuron while monitoring membrane potential fluctuations in the other neurons. While this approach has revealed synaptic connectivity in the neocortex (Karnani et al., 2016a, 2016b), we found no connections in LH (0 connected in 248 tested putative connections, Figure 1C).

**Fig. 1.**
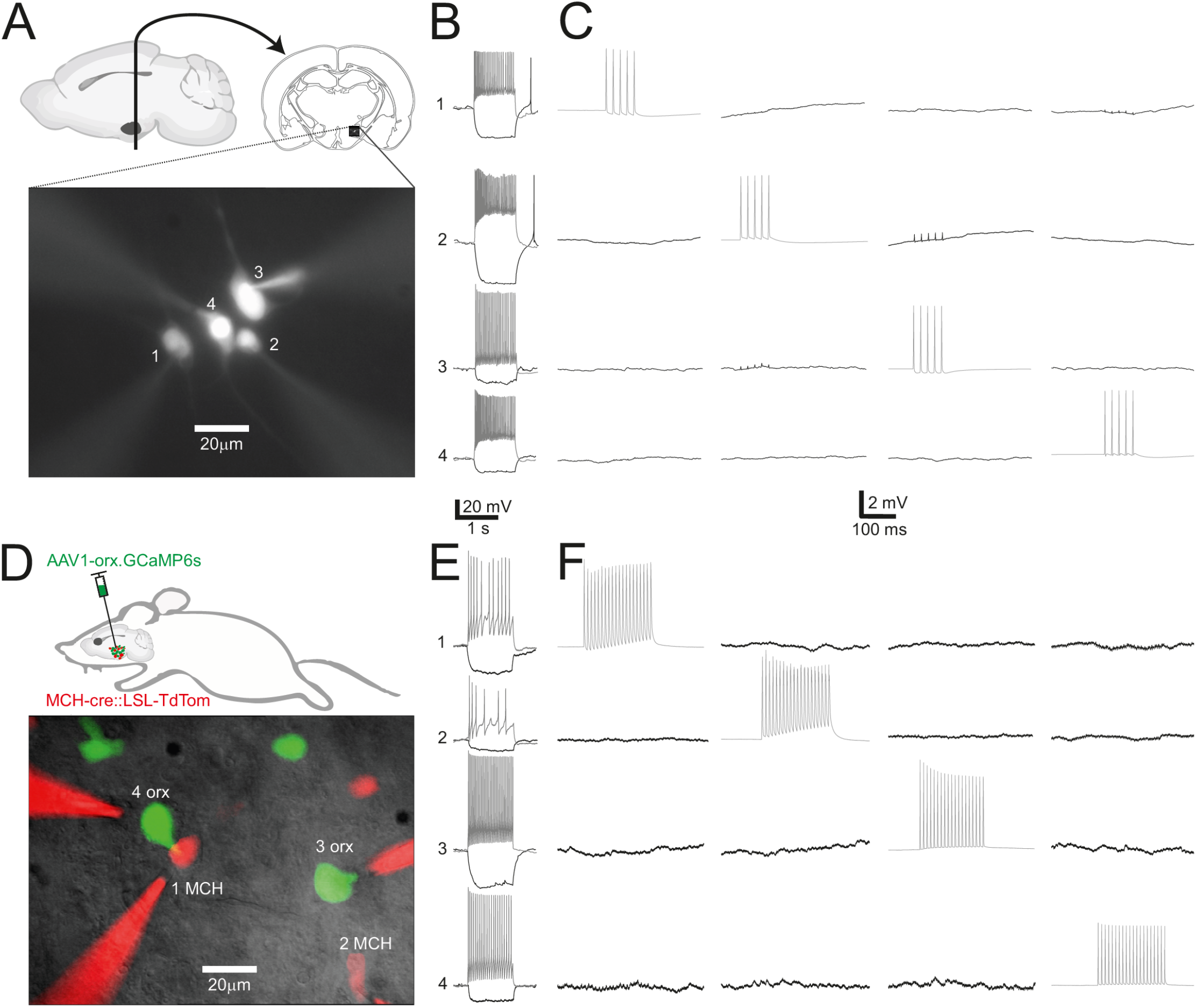
Example quadruple whole cell recordings from lateral hypothalamic slices. A, Schematic of recording location and fluorescence micrograph of a recording configuration of four unidentified (no-marker) neurons in LH. B, membrane potential and firing responses of neurons shown in A to a −30pA (black) and 150pA (gray) current injection. Cells were held at −50mV in baseline. C, Example connectivity recording of cells in A and B, with tested presynaptic cells firing 5 action potentials at 50Hz (gray) and tested postsynaptic cells showing lack of synaptic responses (black). Traces are averages of 50 trials. D, Schematic of labelling strategy for identifying orexin and MCH neurons using three transgenes, and fluorescence micrograph of example recording configuration with two MCH cells and two orexin cells. E, neurons in D recorded as in B. F, neurons in E and D recorded as in C, except with a longer train of action potentials. Traces are averages of 30 trials. Scale bars below B and C apply to E and F respectively. Vertical scale in C and F only applies for black traces.

As it is possible that connectivity is specific to neuronal subtypes, we used multiple transgenes to identify genetically defined LH subpopulations in acute slices through fluorescent protein expression. This approach (see Methods) allowed us to target up to two identified populations (e.g., orexin and MCH neurons labelled in Figure 1D,E) as well as the non-marked (n.m.) neurons outside these populations. With this approach we also found zero connectivity in all but one category (1/79 tested GAD65-GFP → vglut2 putative connections was connected), between and within the cardinal LH populations, orexin, MCH, vglut2, GAD65-GFP, GAD65-cre, VGAT-cre and n.m. neurons (Figures 1F and 2A; overall, 1 synaptic connection was found among 2074 tested; Bonnavion et al., 2016; Mickelsen et al., 2019).

**Fig. 2.**
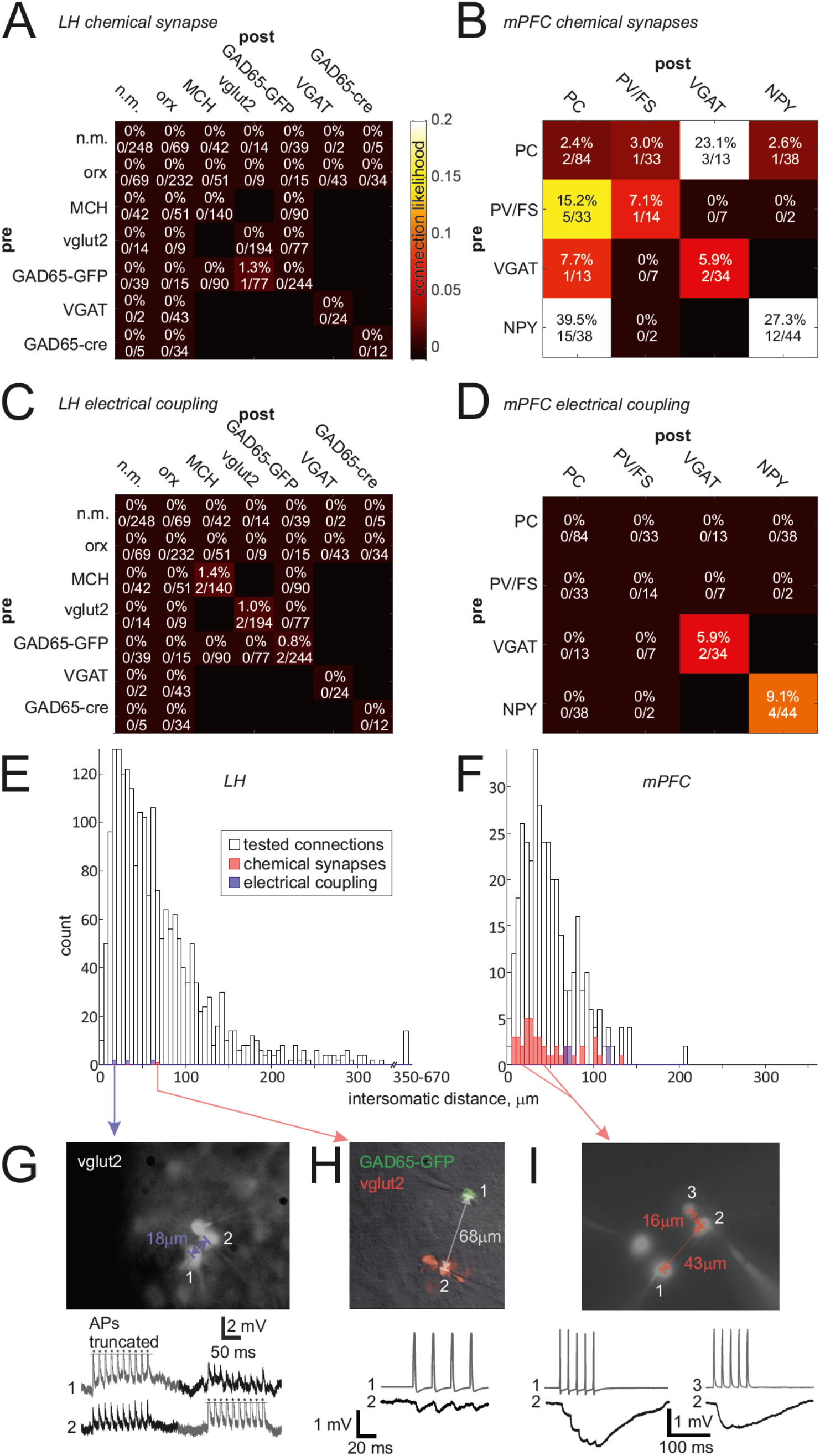
Analysis of connectivity recordings in LH and mPFC. A, overall synaptic connectivity within LH across 2074 tested connections showing 1 synapse. B, overall synaptic connectivity within mPFC across 362 tested connections, showing 43 synapses. C, Electrical coupling in LH was found in 3 cell pairs among 1037 tested. D, Electrical coupling in mPFC was found in 3 cell pairs among 181 tested. E and F, distances between cell somata in tested connections in LH (E) and mPFC (F). G-I, example micrographs and voltage recordings of identified connections. Action potentials (APs) truncated in G, as indicated by asterisks. H shows the only local chemical synapse found within LH. I shows an example recording from mPFC with a NPY neuron (3) and a PV/FS neuron (1) inhibiting a pyramidal cell (2). Vertical scale bars in H and I apply only to postsynaptic traces (black).

We took several measures to control for potential sources of artefact in this important result. We verified that slice geometry is not a confounding factor by testing connections in horizontal (0 connected in 74 tested), sagittal (0 connected in 50 tested) and coronal slices (1 connected in 1950 tested). Because the distance from slice surface can conceivably affect connectivity through limiting the available space where synapses might have existed in the intact brain, we monitored the depths of recorded neurons (37.2 ± 22.5 μm, range 12-123 μm, n = 50). These depths are within the reported range (5-130 μm) of reliable connectivity measurements reported before (Fino and Yuste, 2011). As preservation of connections can vary across individual slices, we report that in the slice with the one recorded synapse, 17 other trials revealed no connections, and in the adjacent slices from that animal, 36 other trials revealed no connections. As age of animals can affect connectivity, we report the age of this animal was P45, which is far from the low end of distribution of ages in this study (P21-201, mean P94.8 ± 42.3). Finally, to show that nothing in the methodology systematically affects synaptic connectivity, we recorded with the same equipment and preparation protocol, a control dataset from medial prefrontal cortex (mPFC, including infralimbic, prelimbic and anterior cingulate cortices). In this control data 43 synaptic connections were observed out of 362 trials (Figure 2B).

These connections were more frequent among inhibitory neurons as has been shown by many other studies (Karnani and Jackson, 2018; Karnani et al., 2014).

Bidirectional electrical coupling among LH neurons was observed on three occasions, once each among pairs of MCH, GAD65-GFP and vglut2 neurons (Figure 2C). This overall electrical coupling rate of 0.3% (6/2074) was contrasted by a rate of 1.7% (6/362) in mPFC (Figure 2D). In both brain structures this very sparse electrical coupling was found within genetically defined subpopulations, which is typical in the cortex (Karnani and Jackson, 2018). Electrical coupling coefficient (see Methods) was 0.075 ± 0.102 in LH and 0.018 ± 0.013 in mPFC (P=0.2), values which are typical for adult CNS (Alcamí and Pereda, 2019).

As synaptic connectivity is known to be distance dependent, we measured intersomatic distances from all experiments (Figure 2E,F). Distances in LH (mean 71.6 ± 65.7 μm) were significantly higher than distances in mPFC (mean 50.3 ± 32.4 μm, P<10^−6^ by Wilcoxon rank sum test), because, failing to find connections at short distances, we searched for possible longer distance connections. The range of covered distances was broader in LH (3.0 – 664.5 μm) than mPFC (4.4 – 208.7 μm), and encompassed all mPFC distance bins and exceeded them in numbers (Figure 2E,F). The intersomatic distance bins of the discovered synapses were well sampled, indicating that the measured connectivity rates are reliable. The discovered synapse in LH was inhibitory and weaker than typical synapses in mPFC (Figure 2G-I). We therefore conclude that local synaptic connectivity in LH, bench-marked to that in neocortex, is nearly non-existent.

### Single synapse equivalent connections with optogenetic population activation

Having found a surprising lack of synaptic connectivity with multiple whole cell recordings, we sought to reconcile our result with previously demonstrated connections using optogenetics (Apergis-Schoute et al., 2015; Ferrari et al., 2018; Kosse and Burdakov, 2019; Kosse et al., 2017). We first assessed several population specific connections optogenetically, confirming in each case that the optogenetically labelled presynaptic population was firing action potentials in response to light (Figure 3A-C). Connections were rarer than ‘non-connections’ (43/136, Figure 3C), and the postsynaptic response amplitudes were low considering they arise from firing of thousands of presynaptic neurons (MCH→orx 0/5; orx→GAD65-GFP 1/16, 0.3 mV; GAD65-cre→orx 2/25, 2.0 and 3.1 mV; VGAT→orx, 2/24, 0.6 and 0.6 mV; orx→n.m. 1/8, 0.1 mV; MCH→n.m. 3/9, 0.3 ± 0.2 mV; GAD65-cre→n.m. 3/8, 3.6 ± 5.4 mV; vglut2→GAD65-GFP 9/15, 1.4 ± 1.6 mV; vglut2→n.m. 22/26, 2.8 ± 2.8 mV). Latencies from light to response onset were consistent with monosynaptic responses (orx→GAD65-GFP 2.5 ms; orx→n.m. 2.4 ms; GAD65-cre→orx 11.5 and 8.2 ms; VGAT→orx, 7.3 and 10.6 ms; orx→n.m. 2.4 ms; MCH→n.m. 10.5 ± 5.0 ms; GAD65-cre→n.m. 5.6 ± 1.5 ms; vglut2→GAD65-GFP 7.6 ± 4.5 ms; vglut2→n.m. 6.4 ± 2.2 ms).

**Fig. 3.**
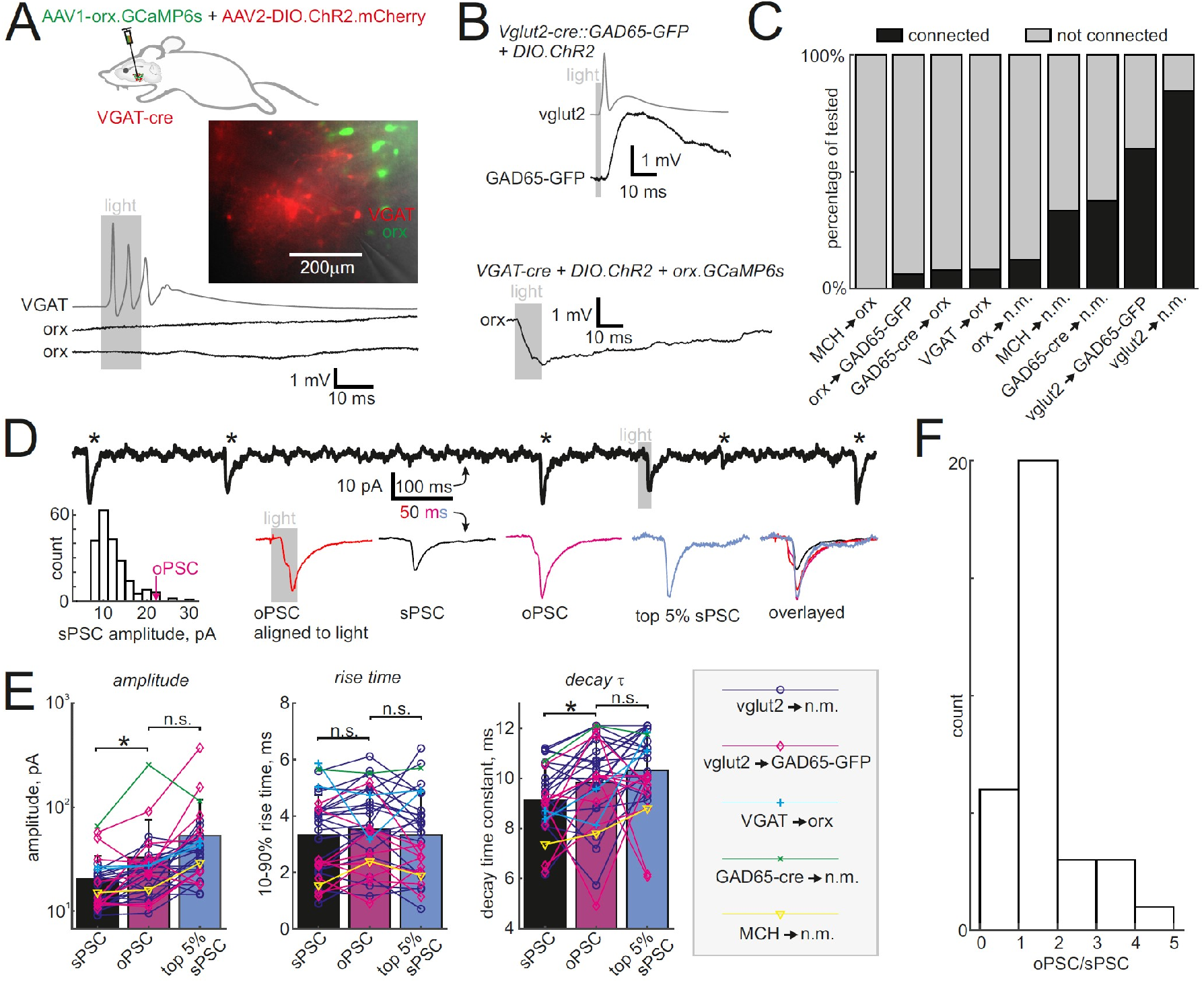
Optogenetic connections within the LH are consistent with ultra-sparse synaptic connectivity. A, Above, schematic of opsin and label delivery strategy to identify connections from VGAT neurons to orexin neurons. Middle, micrograph of recording configuration of a VGAT neuron with channel rhodopsin and two orexin neurons. Below, current clamp recording shows a typical lack of synaptic response in orexin neurons while the simultaneously recorded VGAT neuron spikes in response to the light stimulus. B, Examples of identified connections from the optogenetically activated vglut2 population to a GAD65-GFP neuron (above) and from the VGAT population to an orexin neuron (below). Amplitude scale bars in A and B only apply to black traces. Gray traces illustrate spiking in the channel rhodopsin bearing population. C, Summary data from all recordings showing that overall only 32% (43/136) of recorded neurons had a postsynaptic response when thousands of presynaptic neurons were activated. D, Example voltage clamp (−60 mV) recording of oPSC and sPSCs from the GAD65-GFP neuron shown in B, when vglut2 neurons were optogenetically tagged. Above trace is a segment of the recording showing a light induced PSC and several sPSCs (*). Bottom, from left, histogram of sPSC amplitudes and the average oPSC amplitude labelled, average oPSC across all light stimuli, average sPSC, average oPSC (detected and aligned with same method as the sPSC for a reliable comparison), average of the largest 5% of sPSCs, and the average waveforms overlayed. E, Summary data of average PSCs as labelled in D across all voltage clamp data, showing that oPSCs were larger than sPSCs, but not significantly different from the top 5% of sPSCs. * P < 0.05; n.s. P > 0.05. Y-axis in amplitude plot is logarithmic. F, Histogram of oPSC/sPSC amplitude ratio for all neurons in E, showing the majority of optogenetic connections can be explained by 1-2 sPSCs.

We wondered whether the optogenetically-evoked postsynaptic currents (oPSCs) would be on par with spontaneous synaptic currents (sPSCs) which typically arise from spontaneous discharge of single synapses or individual presynaptic neurons. This comparison, which has never been documented, would offer an explanation wherein the previously demonstrated optogenetic connections may have arisen from an ultra-sparse connectivity coupled with activation of thousands of neurons, with possibly only one of them actually presynaptic to the recorded neuron. We therefore recorded oPSCs and sPSCs in continuous recordings withing the same neurons (Figure 3D). Across cells, oPSCs were 61% bigger than sPSCs (sPSCs 20.3 ± 13.5 pA, oPSCs 32.8 ± 42.7 pA, P<0.05, Figure 3E). However, the average oPSC amplitudes were typically within the distribution of sPSC amplitudes (Figure 3D). Furthermore, oPSCs were not significantly different (−37%, P>0.05) from the average of the 5% largest sPSCs from each cell (top 5% sPSCs 52.8 ± 64.4 pA, oPSCs 32.8 ± 42.7 pA, Figure 3E), suggesting that the oPSCs can indeed arise from firing of one connected presynaptic neuron. Rise time and decay time constants of oPSCs were similarly on par with the top 5% sPSCs (Figure 3E). In order to estimate how many presynaptic neurons might give rise to the oPSCs, we calculated oPSC/sPSC mean amplitude ratios for each cell (using the mean sPSCs rather than top 5%. The average oPSC/sPSC ratio was 1.6 ± 0.9, and their distribution (Figure 3F) suggests that most oPSCs arise from 1-2 presynaptic neurons and even the largest oPSCs arise from 3-5 among thousands of activated neurons. This result is in line with our multiple patch clamp recordings, and reconciles previous results with ultra-sparse connectivity within the LH.

### Lack of locally generated oscillations in LH

Lastly, we asked what is the functional effect of ultra-sparse connectivity in the LH. The dense excitatory and inhibitory connectivity within the neocortex allows it to generate local network oscillations in the 10-80 Hz beta-gamma band (Buzsáki and Wang, 2012). Does lack of dense connectivity in LH mean that it cannot generate local network oscillations? We used the red-shifted opsin C1V1 driven by the ubiquitous neuronal CaMKII-promoter to induce local network activity (Figure 4A,B). Because the dendrites of LH neurons are not consistently spatially organized, we were unable to measure local field potentials, but instead had to use whole-cell recordings to measure rhythmic effects on transmembrane currents (Adesnik and Scanziani, 2010; Karnani et al., 2016a). We used linear light intensity ramps over 5 s to drive network activity so that a possible ‘sweet-spot’ of network activation would not be missed. However we only found oscillations in mPFC even though neuronal firing was reliably recruited in both mPFC and LH (Figure 4C,D; n=10 neurons in each area). To demonstrate that the membrane current oscillation in mPFC neurons was not due to intrinsic properties of the recorded neurons, which may differ from LH neurons, we washed on a cocktail of synaptic blockers which abolished the oscillation (Figure 4E,F, average 10-80 Hz band power induced by light ramp dropped by 99%, P = 10^−4^ by paired t-test, n = 5). On average LH neurons had 11% of the 10-80 Hz band power increase seen in mPFC neurons during the light ramp, and the power in LH neurons was not significantly different from power in mPFC neurons under synaptic blockade (Figure 4F). Therefore, the data suggests that the ultra-sparse connectivity in LH makes it unable to generate local beta and gamma oscillations.

**Fig. 4.**
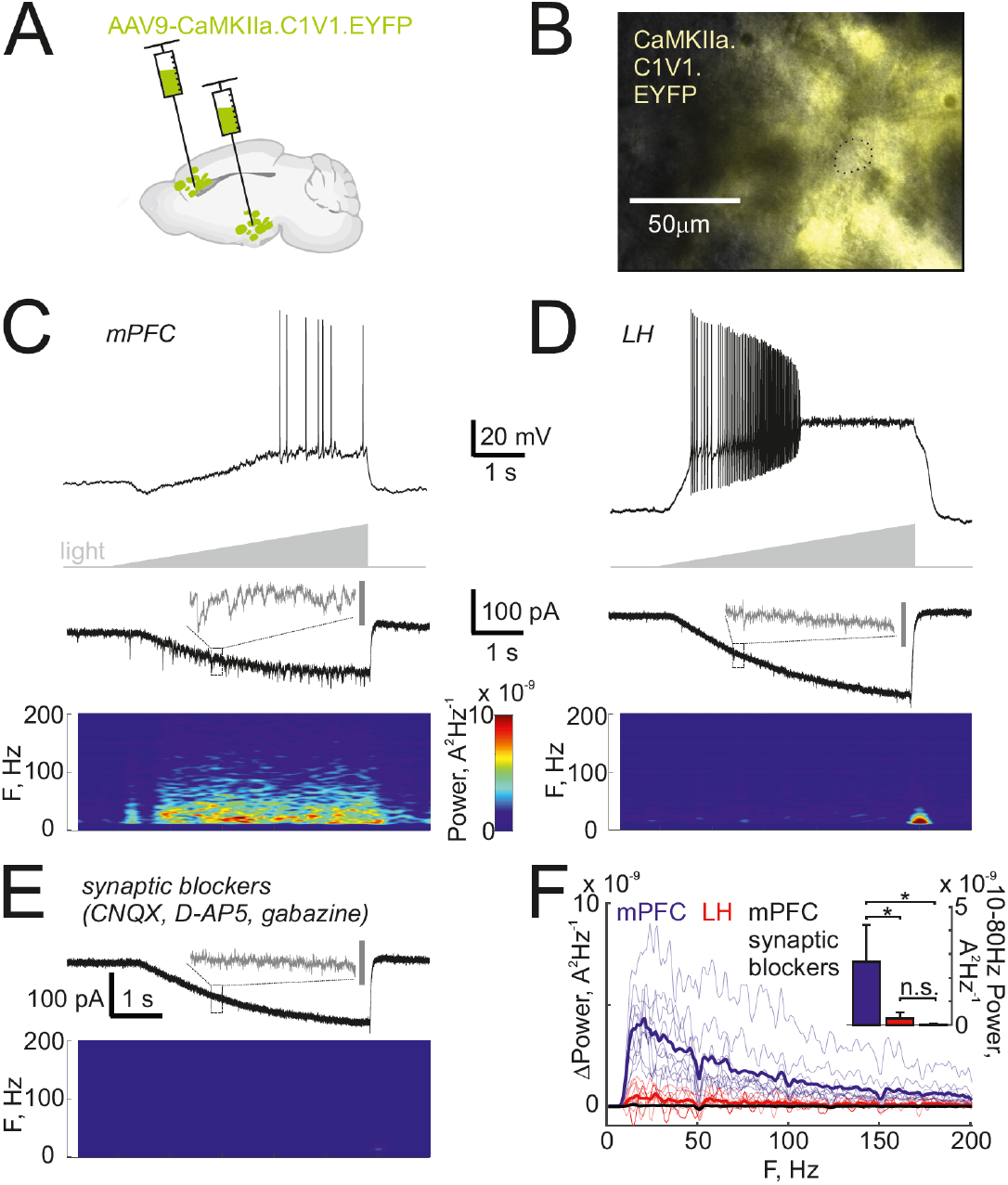
Optogenetically induced local network oscillations are absent in LH. A, Schematic of opsin delivery to LH and mPFC. B, Example epifluorescence image of live LH neurons showing abundant expression of CaMKII.C1V1.EYFPC and the recorded neuron circled by a dashed line. C, Example recordings in mPFC. Above, example of light ramp induced firing in current clamp recording. Middle, voltage clamp recording (−60 mV) during the light ramp-induced synaptically driven oscillatory currents. Bottom, spectrogram showing increasing power in the beta-gamma band (10-80 Hz) during the light ramp. D, Same as C but in LH, showing absence of beta-gamma oscillations. E, Same voltage clamp recording as in C after adding synaptic blockers, showing the oscillation is dependent on local synaptic connectivity. Colormap in C applies to D and E. Time scale is the same in all plots C-E except for gray insets on voltage clamp traces are expanded views of the indicated 200ms segment (dashed box) and the gray scale bar is 50pA (expanded views are scaled identically for comparison). F, Power increase across frequency bands during the light ramp, showing presence of light-driven beta-gamma oscillations in mPFC (recordings from each cell in light blue, average in thick blue, n=10) but not in LH (recordings from each cell in light red, average in thick red, n=10), and deletion of mPFC oscillations by synaptic blockade (recordings from each cell in gray, average in thick black, n=5). Inset shows average power in the 10-80 Hz band across cells. * P < 0.05; n.s. P > 0.05.

## Discussion

We have found with multi-neuron whole cell recordings that the LH does not contain neocortical-like densely connected microcircuits (Figure 1,2). This is in line with the many optogenetically non-connected neurons in this study (Figure 3A,C) and previous studies (Ferrari et al., 2018; Kosse and Burdakov, 2019), as well as the orders of magnitude stronger inputs arising from optogenetically activated long-range sources of input to LH (Giardino et al., 2018; González et al., 2016a). Although we can qualitatively reconcile existing literature on LH intra-connectivity, our findings diverge quantitatively from some published optogenetic datasets: Firstly, we found VGAT-cre → orexin neuron optogenetic connections only in 2/24 cases while Ferrari and colleagues found them in 71/116 cases (Ferrari et al., 2018). This may be explained by the vector used to deliver the opsin to VGAT-cre cells, as Ferrari et al. used AAV5 which is known to have retrograde tropism (Aschauer et al., 2013) and could therefore have expressed in terminals of VGAT-cre neurons projecting into the LH from anywhere in the brain. We used AAV2 which has not been reported to traffic retrogradely. Secondly, we found an orexin → GAD65-GFP neuron optogenetic connection only in 1/16 cases whereas Kosse and colleagues found them in 5/6 cases (Kosse et al., 2017). This may also be explained by opsin delivery strategy as Kosse et al., used a cre-dependent channel rhodopsin injected into an orexin-cre line where ~14% of cells with opsin do not contain orexin (Schöne et al., 2012). We used a virus with an orexin promoter, which delivers the opsin to less cells outside the orexin population, ~4% (Karnani et al., 2020). Such an increase in specificity of the labelled population can explain the difference in effect magnitude. Indeed the inherent difficulty in interpreting full field optogenetic connectivity data lies in the characterization of the presynaptic population, which typically cannot be done in the same slices where the connection was recorded.

The extreme sparseness of LH intraconnectivity has many implications. As the LH does not contain a gamma generator (Figure 4) needed to receive and send gamma rhythmic communications, an information transfer protocol based on gamma coherence (Akam and Kullmann, 2014) is unlikely to be utilized by the LH. Dense connectivity in the neocortex is based on inhibitory interneurons which migrate into the cortical scaffold from the ganglionic eminences during development (Silva et al., 2019). The migrating interneurons do not go into the adjacent hypothalamus due to the presence of nonpermissive factors which directs them into the frontal migratory streams toward neocortex and striatum (Wichterle et al., 2003). Thus it may be that the hypothalamus lacks dense connectivity as a consequence of acting as a ‘sheep-dog’ to migrating interneurons. In line with this, two other subcortical circuits at the rostro-caudal level of the LH lack intrinsic synaptic connectivity, the subthalamic nucleus (Steiner et al., 2019) and thalamic reticular nucleus (Hou et al., 2016). It stands to reason that the denser the microcircuit, the higher its malleability, noise and error-proneness. Therefore dense connectivity may pose a significant risk to survival. As nearby brain regions are thought to have arisen by duplication and elaboration in evolution (Cisek, 2019), one intriguing possibility is that dense local synaptic microcircuits evolved at later stages as a risky evolutionary experiment in structures that are less vital than the hypothalamus. Lastly, the ultra-sparse connectivity in LH implies that any integration and filtering of afferent input to the LH occurs primarily within individual LH neurons. Consequently, coordination of activity in upstream networks is required for the rapid, coordinated activity of LH neurons during behaviour (González et al., 2016a; Karnani et al., 2020).

## Materials and Methods

### Animals

Animal handling and experimentation was approved by the UK government (Home Office) and by Institutional Animal Welfare Ethical Review Panel or carried out according to recommendations in the Animal Welfare Ordinance (TSchV 455.1) of the Swiss Federal Food Safety and Veterinary Office, and were approved by the Zürich Cantonal Veterinary Office. Animals of both sexes, aged 21-180 days at the beginning of the procedures were used and were housed in a controlled environment on a reversed 12h light-dark cycle with food and water ad libitum. Breeders were LSL-TdTom (Ai14), vglut2-cre (Slc17a6^tm2(cre)Lowl^), VGAT-cre (Slc32a1^tm2(cre)Lowl^), GAD65-cre (Gad2^tm2(cre)Zjh^), MCH-cre (Tg(Pmch-cre)1Lowl), GAD65-GFP (Karnani et al., 2013) and WT C57BL6 mice for LH slices and NPY-GFP (Tg(Npy-hrGFP)1Lowl), PV-cre (Pvalb^tm1(cre)Arbr^) or VGAT-cre for mPFC slices, and were obtained originally from the Jackson Laboratory. Several strategies were used to obtain animals with two marker labelled populations in LH as follows. Crossing the breeders to yield vglut2-cre::LSL-TdTom::GAD65-GFP, MCH-cre::LSL-TdTom::GAD65-GFP, or VGAT-cre::LSL-TdTom, MCH-cre::LSL-TdTom, GAD65-cre::LSL-TdTom injected with an orexin promoter virus (ORX.GCaMP/ORX.C1V1) explained below, or a cre line injected with a mixture of a floxed virus and an orexin promoter virus explained below.

### Virus injections

Mice were injected stereotactically with 100-150nl of AAV1-hORX.GCaMP6s (2.5×10^12^ GC/ml, U Penn vector core), AAV1-hORX.C1V1(t/s).mCherry (>10^13^ GC/ml, Vigene), AAV2-EF1a.DIO.hChR2(E123T/T159C).mCherry (7.3×10^12^ UNC GTC Vector Core), or 300nl of 1:1 mixed AAV1-hORX.GCaMP6s and AAV2-EF1a.DIO.hChR2(E123T/T159C).mCherry, or 400nl of AAV9-CaMKIIa.C1V1(t/t).TS.EYFP (≥10^13^ CG/ml, Addgene). Orexin promoter virus expression specificity has been characterized previously (González et al., 2016b; Karnani et al., 2020). The used viral loads were similar to, or higher than, those used in compared studies (Apergis-Schoute et al., 2015; Ferrari et al., 2018; Kosse and Burdakov, 2019; Kosse et al., 2017). For surgery, mice were anesthetized with isoflurane, the scalp was infiltrated with lidocaine, opened, and a 0.2 mm craniotomy was drilled at 0.9 mm lateral, 1.4 mm posterior from Bregma. A pulled glass injection needle was used to inject virus 5.4 mm deep in the brain at a rate of 50 nl/min. The surgery was repeated similarly in the other hemisphere, except when using AAV9-CaMKIIa.C1V1(t/t).TS.EYFP, in which case the surgery was repeated in the other hemisphere at the PFC coordinates 0.4 mm lateral, 1.7 mm anterior from Bregma, and injection at 1.5 mm depth. After removal of the injection needle, the scalp was sutured and animals received 5 mg/kg carprofen injections for two days as post-operative pain medication. Virally injected animals expressed the proteins for an average of 48 ± 18.4 days before the experiment (range 27-91 days).

### Preparation of acute slices

Coronal brain slices from P21-180 animals were prepared after instant cervical dislocation and decapitation. The brain was rapidly dissected and cooled in continuously gassed (95% O_2_ and 5% CO_2_), icy cutting solution containing (in mM): 90 N-methyl-D-glucamine, 20 HEPES, 110 HCl, 3 KCl, 10 MgCl_2_, 0.5 CaCl_2_, 1.1 NaH_2_PO_4_, 25 NaHCO_3_, 3 pyruvic acid, 10 ascorbic acid and 25 D-glucose. 350 μm thick coronal brain slices were cut on a vibratome (Campden) and allowed to recover for 5-15 min at 37 °C in cutting solution, followed by 45-55 min at 22 °C in artificial cerebrospinal fluid (ACSF) containing (in mM): 126 NaCl, 3 KCl, 2 MgSO_4_, 2 CaCl_2_, 1.1 NaH_2_PO_4_, 26 NaHCO_3_, 0.1 pyruvic acid, 0.5 L-glutamine, 0.4 ascorbic acid and 25 D-glucose, continuously gassed with 95% O_2_ and 5% CO_2_.

### Slice electrophysiology

Patch clamp recordings were performed in a submerged chamber with 3-5 ml/min superfusion with ACSF, continuously gassed with 95% O_2_ and 5% CO_2_. A modified Olympus upright microscope with diascopic gradient contrast optics and episcopic fluorescence was used to identify neurons in slices. 3-7 MOhm patch pipettes were filled with intracellular solution containing (in mM): 130 K-gluconate, 5 NaCl, 2 MgSO_2_, 10 HEPES, 0.1 EGTA, 4 Mg-ATP, 0.4 Na-GTP, 2 pyruvic acid, 0.1 Alexa-594, 0.1% biocytin, and ~10 mM KOH (to set pH to 7.3). Whole cell recordings were not analysed if the access resistance was above 25 MOhm. Recordings were sampled at 10 or 20 kHz and low-pass filtered at 3 kHz with HEKA EPC10 usb amplifiers and acquired with HEKA patchmaster software. Current clamp data were recorded at by injecting a steady current to set membrane potential at −50mV. Action potential trains were elicited with 50Hz, 1ms, 1nA current steps. Voltage clamp data were recorded at a holding voltage of −60mV to study excitatory PSCs and 0mV to study inhibitory PSCs. These protocols and solutions have been used to reliably study cortical microcircuits in this paper (Figure 2) and others (Jackson et al., 2018; Karnani et al., 2016a, 2016b). Electrical coupling coefficient was measured during a current step as the size of postsynaptic cell voltage change divided by size of presynaptic voltage change (Alcamí and Pereda, 2019). Opsins were stimulated with green (~16 mW/mm^2^, for orx-C1V1) or blue (~10 mW/mm^2^, for ChR2) light from a xenon lamp (Sutter lambda 4DG controlled from HEKA patchmaster) through a TRITC-filter or with a 532 nm green laser (Laserglow) for linear light ramps from 0 to ~20 mW/mm^2^. Patch clamp data were analyzed in Matlab. Spectrograms were generated in Matlab with the spectrogram function, using 0.5s windows and 95% overlap, after high-pass filtering at 10Hz. Chemicals for making solutions were purchased from Sigma-Aldrich, except synaptic blockers CNQX, D-AP5 and gabazine which were from Tocris.

### Statistics

All data are shown as mean ± s.d. unless stated otherwise. Statistical significance was determined by paired or unpaired Student t-test or Wilcoxon signed rank test as stated. All statistics were performed using statistical functions in Matlab.

## Author Contributions and Notes

Conceptualization: MMK. Methodology: MMK. Software: MMK. Analysis: MMK. Investigation: MMK. Resources: DB. Writing – original draft preparation: MMK. Writing – review and editing: MMK, DB. Funding acquisition: MMK, DB.

The authors declare no competing interests.

## Acknowledgments

We thank Jesse Jackson for helpful comments and discussion. This project has received funding from the European Union’s Horizon 2020 research and innovation programme under the Marie Skłodowska-Curie grant agreement DRIVOME (grant agreement 701986). This work was funded by The Francis Crick Institute, which receives its core funding from Cancer Research UK, the UK Medical Research Council.

